# What is valued in conservation? A framework to compare ethical perspectives

**DOI:** 10.1101/2020.09.04.282947

**Authors:** Guillaume Latombe, Bernd Lenzner, Anna Schertler, Stefan Dullinger, Michael Glaser, Ivan Jarić, Aníbal Pauchard, John R. U. Wilson, Franz Essl

**Affiliations:** BioInvasions, Global Change, Macroecology – Group, Department of Botany and Biodiversity Research, University Vienna, Rennweg 14, 1030 Vienna, Austria; Institute of Evolutionary Biology, The University of Edinburgh, King’s Buildings, Edinburghn EH9 3FL, United Kingdom; Division of Conservation Biology, Vegetation and Landscape Ecology, Department of Botany and Biodiversity Research, University of Vienna, Rennweg 14, 1030 Vienna, Austria; Biology Centre of the Czech Academy of Sciences, Institute of Hydrobiology, Na Sádkách 702/7, 370 05 České Budějovice, Czech Republic; University of South Bohemia, Faculty of Science, Department of Ecosystem Biology, Branišovska 1645/31a, 370 05 České Budějovice, Czech Republic; Laboratorio de Invasiones Biológicas (LIB), Facultad de Ciencias Forestales, University of Concepcion Victoria 631, Concepción, Chile; Institute of Ecology and Biodiversity (IEB), Santiago, Chile; South African National Biodiversity Institute, Kirstenbosch Research Centre, Claremont 7735, South Africa; Centre for Invasion Biology, Department of Botany and Zoology, Stellenbosch University, Private Bag X1, Matieland 7602, South Africa

**Keywords:** anthropocentrism, biocentrism, ecocentrism, environmental ethics, impact, invasive alien species, moral values, sentientism, speciesism

## Abstract

Perspectives in conservation are based on a variety of value systems. Such differences in how people value nature and its components lead to different evaluations of the morality of conservation goals and approaches, and often underlie disagreements in the formulation and implementation of environmental management policies. Specifically, whether a conservation action (e.g. killing feral cats to reduce predation on bird species threatened with extinction) is viewed as appropriate or not can vary among people with different value systems. Here, we present a conceptual, mathematical framework intended as a tool to systematically explore and clarify core value statements in conservation approaches. Its purpose is to highlight how fundamental differences between these value systems can lead to different prioritizations of available management options and offer a common ground for discourse. The proposed equations decompose the question underlying many controversies around management decisions in conservation: what or who is valued, how, and to what extent? We compare how management decisions would likely be viewed under three different idealised value systems: ecocentric conservation, which aims to preserve biodiversity; new conservation, which considers that biodiversity can only be preserved if it benefits humans; and sentientist conservation, which aims at minimising suffering for sentient beings. We illustrate the utility of the framework by applying it to case studies involving invasive alien species, rewilding, and trophy hunting. By making value systems and their consequences in practice explicit, the framework facilitates debates on contested conservation issues, and complements philosophical discursive approaches about moral reasoning. We believe dissecting the core value statements on which conservation decisions are based will provide an additional tool to understand and address conservation conflicts.

## INTRODUCTION

The consideration of the moral relationship between people and nature and the consequent ethical obligations for conservation is relatively recent in Western culture. Environmental ethics emerged as an academic discipline in the 1970s (Brennan and Lo 2016) and the concepts of values, duty, and animal welfare, are increasingly appreciated in applied ecology and conservation (Dubois et al. 2017, Díaz et al. 2018). These concepts are complex, and the formulation and implementation of environmental management policies is often associated with conflicts between different groups of stakeholders and between people with different values and interests, for example for the management of charismatic alien species (Redpath et al. 2013, Crowley et al. 2017, Jarić et al. 2020). An examination of how value systems could be explicitly accounted for in conservation decisions could offer opportunities for better identifying conflicts, potentially helping to resolve them, and overall improve environmental management.

Value systems consider more or less inclusive communities of moral patients, defined as the elements with intrinsic or inherent value towards which humans, considered here as the community of moral agents, are considered to have obligations (in the following, for simplicity, we refer to the community of moral patients as the moral community; Table 1). Moral communities can include only humans (anthropocentrism), to further incorporate sentient beings (sentientism), living beings (biocentrism), and collectives (such as species and ecosystems; ecocentrism) (Table 1, Figure 1). The definition of moral communities can also be influenced by additional elements (such as spatial elements in the case of nativism), and, at the assessor level, by personal experience. These value systems underlie different sets of explicit or implicit normative postulates, i.e. value statements that make up the basis of an ethic of appropriate attitudes toward other forms of life, which, in turn, can form the basis of different conservation approaches (Soulé 1985; Table 1). If the normative postulates of different value systems diverge (and excluding considerations that moral reasoning, experience, etc., may change one’s value system), conflicts can arise between different groups of stakeholders whose members share common moral values (Crowley et al. 2017). In particular, conservationists who value biodiversity *per se* [as defined initially by Soulé (1985), called hereafter ‘traditional conservation’ (Table 1)] can be at odds with those who value biodiversity based on human welfare and economic aspects [including ‘new conservation’ (Kareiva and Marvier 2012)], or with those based on animal welfare [‘conservation welfare’ (Beausoleil et al. 2018), or, to a certain extent, ‘compassionate conservation’ (Wallach et al. 2018)]. These issues have been heatedly debated in the literature (Kareiva 2014, Soulé 2014, Doak et al. 2015, Driscoll and Watson 2019, Hayward et al. 2019).

**Table 1.** Glossary of terms as they are used for the purposes of this paper.

| Term | Definition |
| --- | --- |
| Anthropocentrism<br>(strong) | Value system that considers humans to be the sole, or primary, holder of moral standing, and therefore the concern of direct moral obligations. Non-human species are considered only to the extent that they affect the statisfaction of felt preference of human individuals (Norton 1984, Rolston 2003, Palmer et al. 2014). |
| Anthropocentrism<br>(weak) | Value theory in which all values are "explained by reference to satisfaction of some felt preference of a human individual or by reference to its bearing upon the ideals which exist as elements in a world view essential to determinations of considered preferences" (Norton 1984). That is, the value of an individual or species is not only exploitative, but incorporates human experience and the non-utilitarian relationship between humans and nature. |
| Anthropomorphism | “The attribution of human personality or characteristics to something non-human, like an animal, object, etc.” (Oxford English Dictionary n.d.). |
| Biocentrism | Value system considering all living beings as the concern of direct moral obligations (Rolston 2003, Palmer et al. 2014). |
| Collectivism | Value system in which a group or collective has a higher value than the individuals that compose it (Wallach et al. 2018). |
| Compassionate<br>conservation | Conservation approach inspired by virtue ethics based on four tenets: i) do no harm; ii) individuals matter; iii) inclusivity (the value of an individual is independent from the context of the population, e.g. nativity, rarity, etc.); and iv) peaceful coexistence (Ramp and Bekoff 2015, Wallach et al. 2018). |
| Community of<br>moral agents | The group of beings considered to have moral responsibility in their actions (Talbert 2019). We consider it here to be restricted to humans. |
| Community of moral patients | The group of beings considered to have intrinsic moral value, and towards which moral agents have moral obligations (Warren 2000). The size of the group (referred to as the moral community in this work, for simplification) depends on the value system. For example, the moral community is restricted to humans in case of Anthropocentrism. |
| Conservation welfare | Conservation approach aiming at minimizing animal suffering (Beausoleil et al. 2018). |
| Consequentialism | “An ethical doctrine which holds that the morality of an action is to be judged solely by its consequences” (Oxford English Dictionary n.d.). |
| Convergence hypothesis | “If the interests of the human species interpenetrate those of the living Earth, then it follows that anthropocentric and non-anthropocentric policies will converge in the indefinite future” (Norton 1986). |
| Deontology | A normative ethical theory considering that “choices are morally required, forbidden, or permitted” (Alexander and Moore 2016). |
| Ecocentrism | Value system considering that species, their assemblages and their functions, as well as more broadly ecosystems, rather than individuals, are the concern of direct moral obligations (Rolston 2003, Palmer et al. 2014). |
| Empathy | “The quality or power of projecting one's personality into or mentally identifying oneself with an object of contemplation, and so fully understanding or appreciating it.” (Oxford English Dictionary n.d.). Empathy will influence the inherent value given to individuals from other species. |
| Impact (for the purposes of the framework, Eq.1) | Impact refers to any effect that modifies the wellbeing, health or resilience (for non-sentient beings) of an individual, from physical pain to emotional suffering and death (these notions being interrelated, but not equivalent). |
| Inherent value (our definition) | Value possessed by an individual or collective, accounting for their intrinsic value (see definition below) and the effects of multiple context-dependent factors (e.g. charisma, anthropomorphism, organismic complexity, neoteny, cultural importance, religion, or parochialism). For |
|  | example, wolves and dogs may be considered to have similar intrinsic value under sentientism because they have similar cognitive abilities, but may be valued differently by people who own dogs as pets (i.e. due to parochialism). |
| Intrinsic value | Value possessed by an individual or collective as defined by a system of moral valuation, such as anthropocentrism, sentientism, biocentrism or ecocentrism. Once a criteria has been selected in accordance with the system of values (e.g. cognitive ability under sentientism, the choice of a criteria itself may be subjective), intrinsic value is determined by this criteria and context-independent. |
| Invasive alien species | “Plants, animals, pathogens and other organisms that are non-native to an ecosystem, and which may cause economic or environmental harm or adversely affect human health” (*Regulation (EU) No 1143/2014 of the European Parliament and of the Council of 22 October 2014 on the prevention and management of the introduction and spread of invasive alien species*). |
| Moral community | See “Community of moral patients”. |
| Moral dilemma | Situation in which a moral agent regards themselves as having moral reasons to do different, incompatible actions (McConnell 2018). |
| Nativism | Value system considering that species that have evolved in a given location have a higher value in this location than species that have evolved somewhere else. In nativism, value varies spatially (Wallach et al. 2018). |
| Nature despite people | Management conceptual approach aiming at conserving biological diversity (focusing on species and habitats) specifically in response to human impacts on the environment, e.g. sustainable use (Mace 2014). |
| Nature for itself | Management conceptual approach aiming at conserving biological diversity (focusing on wilderness and natural habitats) through human exclusion, for example through the creation of parks and protected areas (Mace 2014). |
| Nature for people | Management conceptual approach aiming at conserving the components |
|  | of nature beneficial to humans (focusing on ecosystems and their services) (Mace 2014). |
| Neoteny | “The retention of juvenile characteristics in a (sexually) mature organism” (Oxford English Dictionary n.d.). |
| New conservation | Discipline aiming at preserving biological diversity through the conservation of natural elements providing services and contribution to human wellbeing (Kareiva and Marvier 2012, Kareiva 2014). |
| Normative postulate | Value statements that make up the basis of an ethic of appropriate attitudes toward other forms of life (Soulé 1985). |
| Parochialism | Ideology in which moral regard is directed “towards socially closer and structurally tighter targets, relative to socially more distant and structurally looser targets”, and, by extension, to species phylogenetically, cognitively, or in appearance closer to humans (Waytz et al. 2019). |
| People and nature | Management conceptual approach considering that humans and nature are interdependent and therefore aiming at achieving compromises in the conservation of nature and human wellbeing (Mace 2014). |
| Relational value | “Preferences, principles, and virtues associated with relationships, both interpersonal and as articulated by policies and social norms [...] Relational values are not present in things but derivative of relationships and responsibilities to them.” (Chan et al. 2016). |
| Sentience | The ability to experience phenomenal consciousness, i.e. the qualitative, subjective, experiential, or phenomenological aspects of conscious experience, rather than just the experience of pain and pleasure (Allen and Trestman 2017). |
| Sentientism | Value system considering sentient beings as the concern of direct moral obligations (Rolston 2003, Palmer et al. 2014). |
| Speciesism | Value system in which some species are considered to have a higher value than others, for various possible reasons (Singer 2009). Speciesism is often used to refer to the superiority of humans, which is a specific expression of speciesism as considered in this paper. |
| Suffering | Negative emotion, sometimes called emotional distress, experienced by sentient beings, and which can result from different causes, including but not limited to physical pain (Dawkins 2008, Farah 2008). |
| Traditional conservation | Discipline aiming at preserving biological diversity through the management of nature, and based on four value-driven normative postulates: “diversity of organisms is good,” “ecological complexity is good,” “evolution is good,” and “biotic diversity has intrinsic value” (Soulé 1985). Traditional conservation is rooted in ecocentrism. |
| Utilitarian value | Value given to an individual or collective by humans, based on its utility. |
| Virtue ethics | Ethical doctrine that emphasizes the virtues, or moral character as the reason for action (Hursthouse and Pettigrove 2018). |

**Figure 1.**
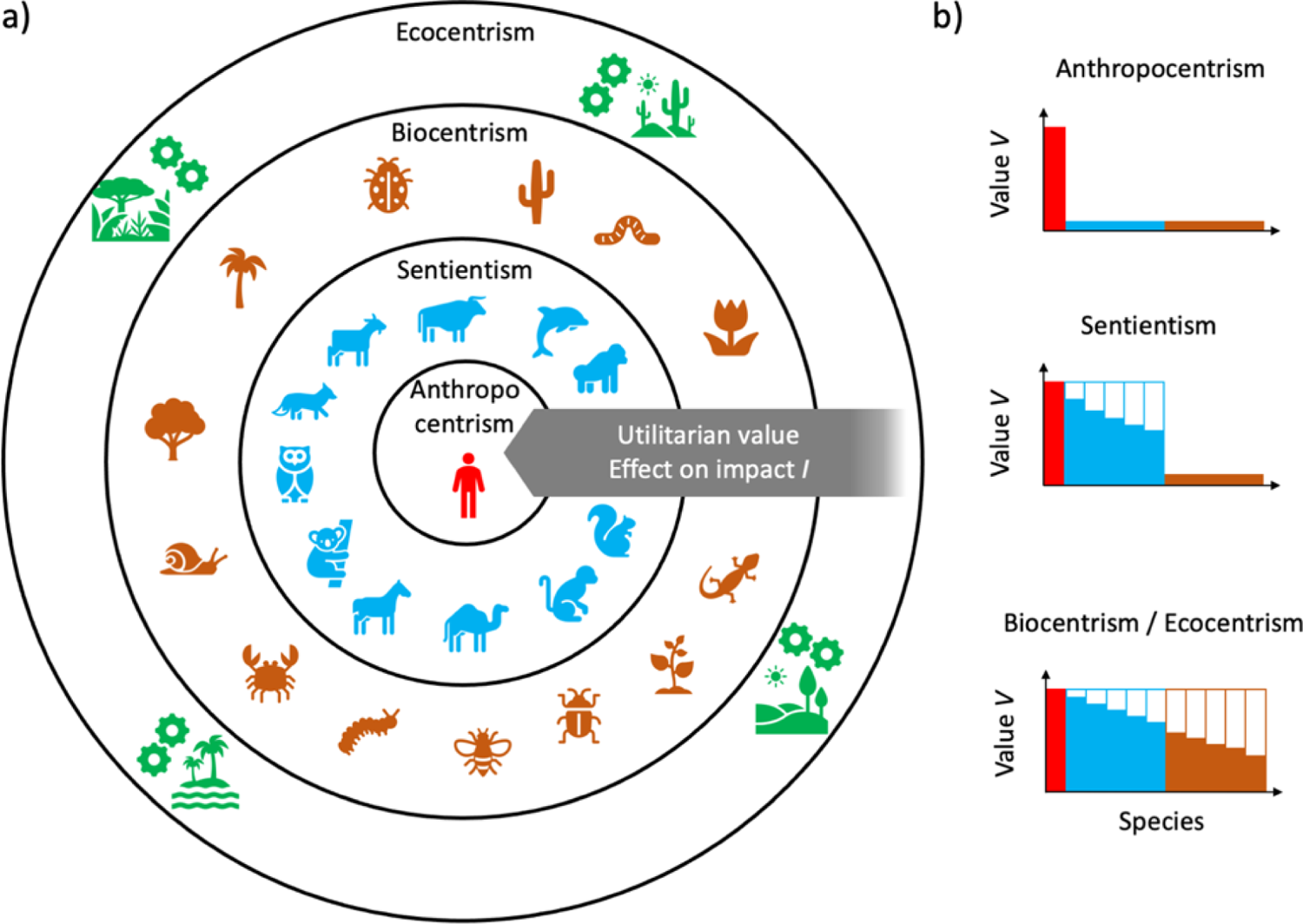
Differences between the moral communities considered by value systems influenced by anthropocentrism, sentientism, biocentrism and ecocentrism (depicted by the nested circles and colours) and how values can differ between members of the different moral communities. a) Anthropocentrism, sentientism and biocentrism all value individuals intrinsically, but consider different moral communities, i.e. their values depend on the category of species they belong to, with {humans} ∈ {sentient beings} ∈ {all living organisms}. Species outside of the moral community may have a utilitarian value for species in the moral community (represented by the arrow), which will be reflected by changes in the impact variable. b) The intrinsic value, in combination with contextual factors, defines the inherent value *V* of an individual or species and the distribution of *V* will change depending on the set of species included in the moral community. Anthropocentrism, sentientism and biocentrism value individuals from different groups of species. Biocentrism and ecocentrism give value to the same group of species, i.e. all living organisms, but while biocentrism values individuals, ecocentrism values ecological collectives, i.e. species or species assemblages and ecosystems. Note that species can have both an inherent and a utilitarian value. Within the moral community, species may have equal inherent values, but subjective perceptions and different value systems may also assign different values to different species. The skewness of the value distribution then indicates the degree or strength of speciesism with respect to the species of reference, assumed here to be the human species, and is influenced by many factors, including charisma, cultural context, etc.

In the following, our aim is to conceptualize and decompose value systems in an explicit, and potentially (but not necessarily) quantifiable, fashion using a common mathematical framework, and to explore their repercussions for the perception of conservation management actions by stakeholders with different value systems. We argue that doing so allows for explicit comparison between these perceptions to identify sources of potential conflicts. First, we recapitulate four archetypal value systems in environmental affairs and relate them to different conservation philosophies. Since identifying commonalities in the perspectives of different parties is key in conflict management (Redpath et al. 2013), we then introduce a formal framework to conceptualise these value systems, and examine how it can be applied to clarify different perspectives. Finally, we discuss opportunities for identifying commonalities between different value systems that may enable identifying widely acceptable solutions to otherwise polarising issues.

### VALUE SYSTEMS AND CONSERVATION PRACTICES

Here, we focus on a Western perspective of value systems that have been internationally considered for environmental policies and the management of nature (Mace 2014). The archetypes of value systems and of conservation approaches were chosen for their importance in the past and present literature and their clear differences, to illustrate our framework. We acknowledge this is a small part of the global diversity of value systems. It would be interesting to see if our framework could be applied to other contexts, to identify its limitations.

### From the valuation of humans to that of ecosystems: a spectrum of value systems in conservation

The Western perspective of moral valuation encompasses a diverse set of value systems with respect to the components of nature that form the moral community. Traditionally, one can distinguish at least four archetypal value systems: anthropocentrism, sentientism, biocentrism, and ecocentrism (Rolston 2003, Palmer et al. 2014) (Table 1; Figure 1).

Anthropocentrism values nature by the benefits it brings to people through ecosystem services, which encompasses economic, biological, and cultural benefits humans can derive from nature (Díaz et al. 2018). One justification for anthropocentrism is that humans are (arguably) the only self-reflective moral beings, and people are both the subject and object of ethics (Rolston 2003), therefore constituting the moral community. In an anthropocentric system, individuals from non-human species only have value based on their benefits or disservices for humans (instrumental or non-instrumental).

Sentientism considers that humans and all sentient animals value their life, and experience pleasure, pain, and suffering. All sentient individuals should therefore also be part of the moral community (i.e. have an intrinsic value). In this view, it is the sentience [e.g. measured through cognitive ability, (Singer 2009)], rather than species themselves, that has intrinsic value.

Biocentrism considers that life has intrinsic value. Although different perspectives on why life has value exist (Taylor 2011), all living organisms are valued equally for being alive, and not differently based on any other trait.

Some ecocentric, or holistic, value systems consider that ecological collectives, such as species or ecosystems, have intrinsic value, independently from the individuals that comprise them. Species can have different values, i.e. speciesism (Table 1), and these values can be influenced by a multitude of factors, discussed in more detail below.

### Subjective elements in the valuation of nature

In practice, the separation between anthropocentrism, sentientism, biocentrism, and ecocentrism is blurry, and values given to different species may vary under the same general approach. For example, biocentrism can range from complete egalitarianism between organisms, i.e. universalism (Table 1), to a gradual valuation resembling sentientism. These four value systems can also interact with other systems that use other criteria than the intrinsic characteristics of individuals to define the moral community. For example, nativism is a system that values organisms indigenous to a spatial location or an ecosystem over those that have been anthropogenically introduced. Nativism can therefore interact with any of the four systems presented above to alter the value attributed to a species in a given context. Finally, the attribution of values to individuals from different species can be deeply embedded in the individual psychologies of the assessor (Palmer et al. 2014, Waytz et al. 2019). Values and personal interests interact in making and expressing moral judgements (Essl et al. 2017). Thus, the archetypes of value systems presented above are rarely expressed in a clear and obvious fashion. Nonetheless, by formalising the archetypes, a framework can be created within which the consequence of conservation actions explored.

To account for the different elements that can be combined to create the concept of value, in the following, we distinguish between ‘intrinsic’, ‘inherent’, and ‘utilitarian’, value (our definitions; Table 1). Intrinsic value is the value possessed by an individual or collective as defined by one of the systems above, and is therefore independent of context. Intrinsic value is based on objective criteria such as cognitive ability. The choice of a criteria may be subjective, but the value is independent of the assessor once the criteria has been defined. This has been termed “objective intrinsic value” by others (Sandler 2012). Inherent value is the value of an individual, species or ecosystem that results from the combination of its intrinsic value and context-specific and subjective factors (note that other scholars have used ‘inherent’ differently, e.g. (Taylor 1987, Regan 2004); here it corresponds to what has also been termed “subjective intrinsic value”; (Sandler 2012)). These factors include charisma (Courchamp et al. 2018, Jarić et al. 2020), anthropomorphism (Tam et al. 2013; Table 1), organismic complexity (Proença et al. 2008), neoteny (Stokes 2007; Table 1), cultural importance (Garibaldi and Turner 2004), religion (Bhagwat et al. 2011), parochialism (Waytz et al. 2019; Table 1), and more generally the relationship between humans and elements of nature (Chan et al. 2016). For example, dogs and wolves may be considered to have similar cognitive abilities objectively, and therefore a similar intrinsic value under sentientism, but dogs may have a higher inherent value for some people because they are in close contact with individuals from this species, i.e. parochialism. Some alien species that did not have any inherent value prior to their introduction have been incorporated in local cultures, therefore providing them a novel and higher inherent value such as horses being linked to a strong local cultural identity in some parts of the USA (Rikoon 2006). Inherent value can often be considered to be fixed at the time scale of a management action, but can nonetheless vary over short time scales in some situations (see the example of the Oostvaardersplassen nature reserve below). Utilitarian value is determined only from an anthropocentric perspective. It is context-dependent and can change rapidly, for example in the case of commercial exploitation.

### Conservation management derived from value systems

Conservation practices can historically be divided into three main categories, closely related to specific systems of moral valuation (Mace 2014). At one extreme, a ‘nature for itself’ (Table 1) view mostly excludes humans from the assessment of the efficacy of conservation management actions (Figure 2). This ecocentric perspective is the foundation of traditional conservation as defined by Soulé (1985), and relies on the four following normative postulates: “diversity of organisms is good,” “ecological complexity is good,” “evolution is good,” and “biotic diversity has intrinsic value” (Soulé 1985). It historically underlies widely-used conservation tools, like the IUCN Red List of Threatened Species (IUCN 2019), in which threat categories are defined in terms of probability of extinction (Mace and Lande 1991) (i.e. a species-level criterion aimed at preserving biodiversity). Ecocentrism is often not limited to the valuation of species, but can encompass wider collectives, i.e. assemblages of species and functions, or ecosystems. This other perspective is captured, for example, by the IUCN Red List of Ecosystems (IUCN-CEM 2016), and it is strongly reflected in international conservation agreements such as the Convention on Biological Diversity (UNEP CBD 2010). In the following we refer to traditional conservation as an ecocentric value system where species are intrinsically valuable (nature for itself; Figure 1) and humans are mostly excluded from management. We acknowledge that this is an archetypal view of traditional conservation, which is used here simply for illustrative purposes.

**Figure 2.**
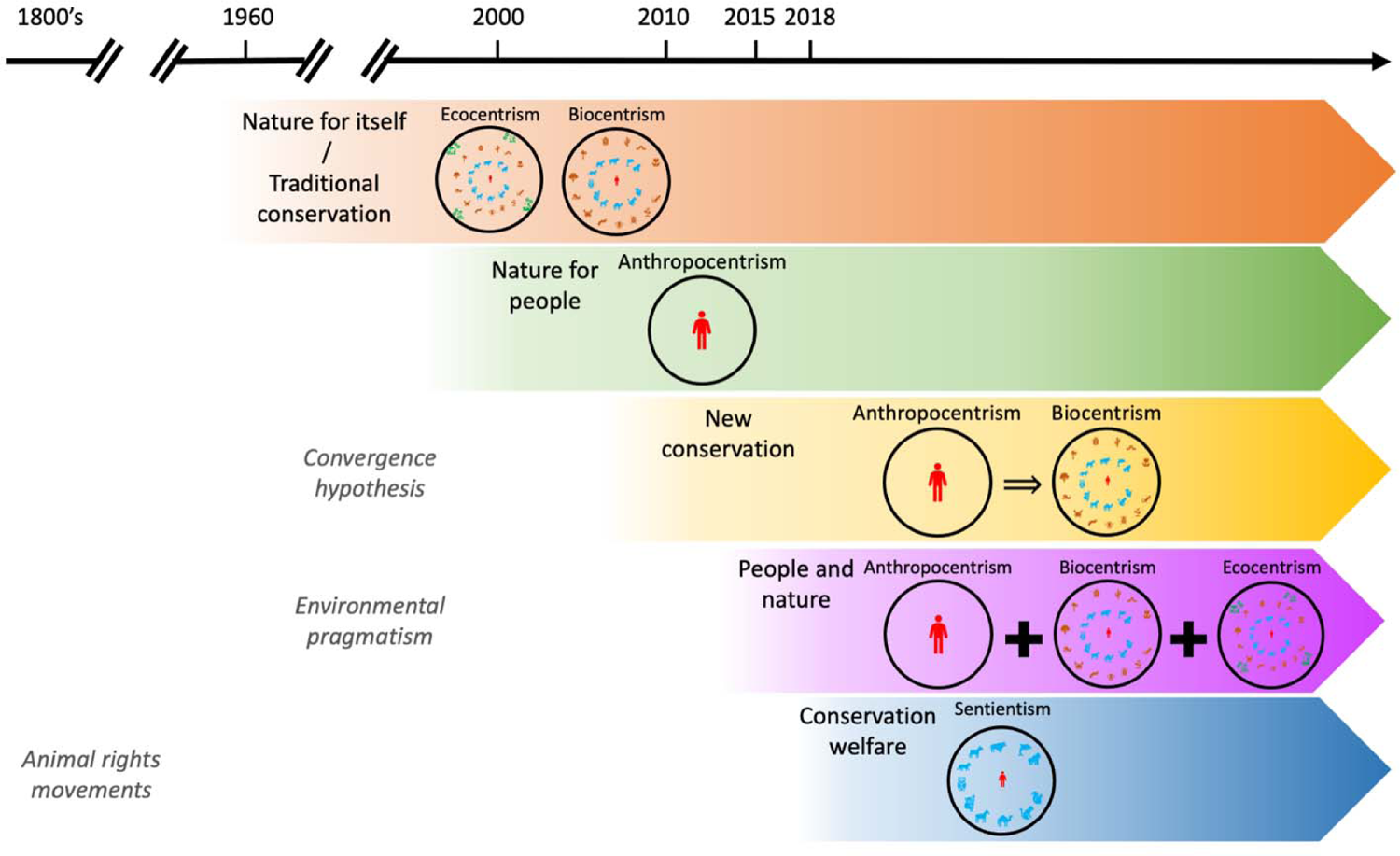
Different value systems (or combination of) correspond to different conservation perspectives, which were introduced at different points in time (the timeline is approximate for illustrative purpose; see also Mace 2014). A nature for itself perspective can be either ecocentric, biocentric, or both. Under new conservation, an anthropocentric perspective is considered necessary to achieve a desirable outcome under a biocentric perspective (=>). Under the people and nature approach, anthropocentric, biocentric and ecocentric perspectives are considered simultaneously (**+**). Underlying concepts and movements pre-dating conservation approaches are indicated in grey italic at the approximate period they originated.

By contrast the more recent, anthropocentric ‘nature for people’ perspective (Mace 2014) values species and ecosystems only to the extent that they contribute to the wellbeing of humans (Figure 2). These values encompass ecosystem services that help sustain human life (Bolund and Hunhammar 1999) or economic assets (Fisher et al. 2008), and can rely on the assessment of species and ecosystem services in terms of their economic value (Costanza et al. 1997), which can be considered as the most general form of utilitarian value, and has also been termed economism (Norton 2000). The ‘nature for people’ perspective can nonetheless incorporate additional measures linked to human wellbeing, such as poverty alleviation or political participation. This more holistic measure of impacts on humans is exemplified by ‘new conservation’, also termed ‘social conservation’ (Miller et al. 2011, Kareiva 2014, Doak et al. 2015) (Table 1; Figure 2). It has been argued that such an anthropocentric perspective will, by extension, help and even be necessary to maximize the conservation of nature (Kareiva and Marvier 2012). Although New Conservation was introduced relatively recently (Figure 2), it follows an older perspective termed the convergence hypothesis, which argues that if human interests depends on the elements of nature, conservation approaches motivated by anthropogenic instrumental or non-anthropogenic intrinsic values should be the same (Norton 1986; Table 1). It is important to note that the exact set of normative postulates proposed by the proponents of new conservation is not clearly defined (Miller et al. 2011), leading to differences of interpretation and heated debates in recent years (Kareiva and Marvier 2012, Kareiva 2014, Soulé 2014, Doak et al. 2015).

More recently, the necessity to account for the interdependence between the health of nature and human wellbeing [i.e. ‘people and nature’ (Mace 2014); Figure 2] has been advocated in the United Nations Sustainable Development Goals (Weitz et al. 2018). This approach lies on the notion of weak anthropocentrism, introduced by the environmental pragmatism movement (Norton 1984, Katz and Light 2013), in which the value of elements of the environment is not only utilitarian, but defined by the relationship between humans and nature (Chan et al. 2016), and therefore is influenced by context and people’s experience (see also the notion of inherent value described above). Similarly, “nature-based solutions” is an approach endorsed by the IUCN, which aims at protecting, sustainably managing, and restoring, natural or modified ecosystems, to simultaneously provide human wellbeing and biodiversity benefits (Cohen-Shacham et al. 2016). The ‘One Health’ approach, endorsed by the Food and Agriculture Organization, the World Health Organization, and the World Organisation for Animal Health also acknowledges the interdependence between the state of ecosystems, human health, and zoonoses (Gibbs 2014). The difference between people and nature and new conservation approaches therefore lies in the fact that it merges anthropocentric and ecocentric systems, rather than considering that the latter will be addressed by focusing on the former (see Section “Nature despite/for/and people” below for details).

Finally, although the animal rights movement, based on sentientism, originated in the 19^th^ century (Salt 1894), it has not, to our knowledge, been formally considered in conservation approaches until recently. Two main approaches can be found in the literature. Conservation welfare (Beausoleil et al. 2018) is a consequentialist perspective that considers conservation under the prism of animal welfare maximisation (Figure 2). Compassionate conservation (Ramp and Bekoff 2015, Wallach et al. 2018), also incorporates animal sentience, but from a virtue ethics perspective. Although conservation welfare aims at aligning with more traditional conservation approaches presented above (Beausoleil et al. 2018), compassionate conservation appears to be set on different values and proposes, for example, to incorporate emotion to provide insight in conservation (Batavia et al. 2021).

### FRAMING MORAL VALUES FOR OBJECTIVE-DRIVEN CONSERVATION

#### Formulation of a mathematical framework

Many of the conflicts in conservation are grounded in the failure to identify and formalize differences in world views, which contain elements of the four archetypes presented above, influenced by cultural norms, economic incentives etc. (Essl et al. 2017). Here, we propose a mathematical formulation as a method to clarify moral discourses in conservation, based on a consequentialist perspective. We therefore consider an objective-driven type of conservation. Our purpose is not to argue about the relevance of consequentialism vs. deontology, or on the place of virtue ethics in conservation. Rather, we consider that, from a management perspective, conservation necessarily includes objective-driven considerations. A better understanding of how and why objectives can differ between stakeholders as a result of their value systems is therefore useful to anticipate and manage potential conflicts. Although some participants of the discourse will be more receptive to discursive than mathematical conceptualisation, we argue that defining concepts as mathematical terms can make differences in value systems and their normative postulates more explicit and transparent, which will be beneficial when used with appropriate stakeholders, even when these terms would be hard to quantify in real life. A mathematical formulation can be seen as a logic way to express relationships between different elements. Doing so can help to identify and facilitate the discussion of shared values and incompatibilities between different environmental policies and management options (Miller et al. 2011), and contribute to manage conflicts (Redpath et al. 2013). In a similar vein, Parker et al. (1999) proposed a mathematical framework for assessing the environmental impacts of alien species. This work was highly influential in the conceptualisation of biological invasions (being cited over 2,000 times until April 2021 according to Google Scholar), rather than by its direct quantitative application. We also acknowledge that this approach has specific limitations, which are discussed below.

Our mathematical formalisation conceptualises the consequences of environmental management actions. As we develop below, these consequences will be defined differently depending on the value system, but can be understood generally as the consequences for the members of the moral community. Under anthropocentrism, these will be consequences for humans; under sentientism, these will be consequences for sentient individuals; under biocentrism and ecocentrism, these will be consequences for biodiversity. We argue that our mathematical formalisation can account for these different value systems (see Appendix S1 for an extension to ecocentrism beyond species and considering wider collectives, i.e. ecosystems), while also accounting for cultural and personal contexts. These consequences *C* can be conceptualised as a combination of the impact of an action on the different species or individuals involved and the value given to said species and individuals under different value systems as follows:

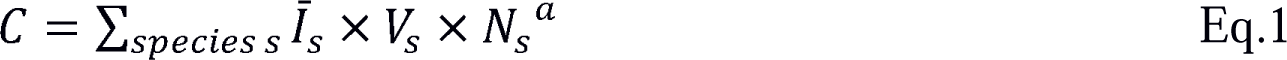

where *I^-^_s_* is a function (e.g. mean, maximum, etc.) of the impact (direct and indirect) resulting from the management action on all individuals of species *s*, *V_s_* is the inherent value attributed to an individual of species *s* (as described above), *N_s_* is the abundance of species *s*, and *a* determines the importance given to a species based on its abundance or rarity (and enables to account for the importance of a species rather than an individual, see below). The unit of *C* depends on how other parameters are defined, which themselves depend on the value system considered. In summary, the higher the impact on species with high values, the higher the consequences.

Inherent value *V_s_* can have a monetary unit or be unit-less depending on how it is defined. It can be continuous or categorical (e.g. null, low, high – quantifiable as 0, 1, 2 or any other quantitative scale). Our definition of inherent value here is extremely broad, as the purpose of this work is not to define what such value should be, rather, it is to be flexible enough to encompass multiple perspectives and the subjectivity of the assessor, and be based on intrinsic, utilitarian or relational values (Chan et al. 2016; Table 1).

The parameter *a* can take both positive and negative values. A value of 1 means that consequences are computed over individuals. If all values *V_s_* were the same, *a* = 1 implies that all individuals in the moral community (Table 1) weigh the same when computing C, irrespective of the species they belong to. This is typical of individual-centred value systems, i.e. sentientism, and biocentrism, whose characteristics (sentience and life) are defined at the individual level. As a result, impacts on larger populations would weigh more on the consequences. As *a* decreases towards 0, the correlation between the value of a species and its abundance decreases. For *a* = 0, the consequence of a management action becomes abundance-independent. For *a* < 0, rare species would be valued higher than common species (or the same impact would be considered to be higher for rare species), for example due to the higher risk of extinction. And for *a* > 1, disproportionate weight is given to abundant species, which are often important for providing ecosystem services (Gaston 2010).

The impact *I_s_* is computed at the individual level. It can be limited to the probability of death of individuals or changes in per capita recruitment rate, thus allowing to compute a proxy for extinction risk if *a* ≤ 0, but can also include animal welfare, biophysical states, etc. As for *V_s_*, continuous or categorical scales may be used. Different measures of impact can be considered under a same system of value, in which case Equation 1 should be applied to each one separately (see section “Application of the mathematical framework” below for details). *I_s_* can only encompass the direct impact of a management action (in a narrow view that only the direct impact of humans, i.e. the moral agents, should be considered, and that the direct impacts from non-moral agents should not be considered), but also include its indirect impact resulting from biotic interactions (considering that, in the context of management and therefore human actions, these indirect impacts are ultimately the result of the actions of the moral agents). One would therefore need to define a baseline corresponding to either i) the lowest possible measurable level of impact (e.g. being alive if death is the only measure of impact, or no sign of disease and starvation for biophysical states; this would obviously be more complicated for welfare), so that *I* would only be positive; ii) the current state of the system, in which case impacts could be positive or negative for different species; or iii) the past state of a system, for example prior to the introduction of alien species (see (Rohwer and Marris 2021) for a discussion on the notion of ecosystem integrity). The duration over which to measure such impact should also be determined. The exact quantification of impact will be influenced by different value systems and personal subjectivity. Some impacts may be considered incommensurable (Essl et al. 2017), therefore falling out of the scope of the framework. The average impact *I^-^_s_* over all considered individuals from a species could be used as a measure at the species-level, as different individuals may experience different impacts, if the management action targets only part of a given population. Using the average impact is not without shortcomings though, since it does not account for potential discrepancies in impacts suffered by different individuals in a population.

In other words, to which point do “the needs of the many outweigh the needs of the few” (Littmann 2016)? Other measures such as the maximum impact experienced by individuals, or more complex functions accounting for the variability of impacts and values across individuals of a same species may also be used, to account for potential disproportionate impacts on a subset of the considered individuals. Under anthropocentric perspectives, impacts are influenced by the utilitarian values of species.

#### Application of the mathematical framework

Considering Equation 1 in an operational fashion, the consequences *C* computed from it can be interpreted as a constructed attribute to measure the achievement of objectives in conservation under different value systems (*sensu* Keeney and Gregory 2005). This may be possible for simple systems with few species and clear categories of values and impacts (Figure 3). However, for complex systems, a quantitative evaluation of Equation 1 will be difficult or impossible. For such systems, the purpose of the framework is not to prescribe how such a constructed attribute should be computed, nor to be used directly as a decision analysis tool (i.e. not to be applied directly). To be used in such a fashion, constructed attributes need to be unambiguous, comprehensive, direct, operational, and understandable by the general public (Keeney and Gregory 2005). Because value systems can be complex, meeting all five criteria is necessarily difficult. Instead, Equation 1 should be seen as a guide to ask questions that are relevant if management shall account for different value systems. By **trying** to evaluate Equation 1, one will have to ask such questions in a systematic fashion (Table 2), while understanding how these questions are conceptually linked with each other.

**Figure 3.**
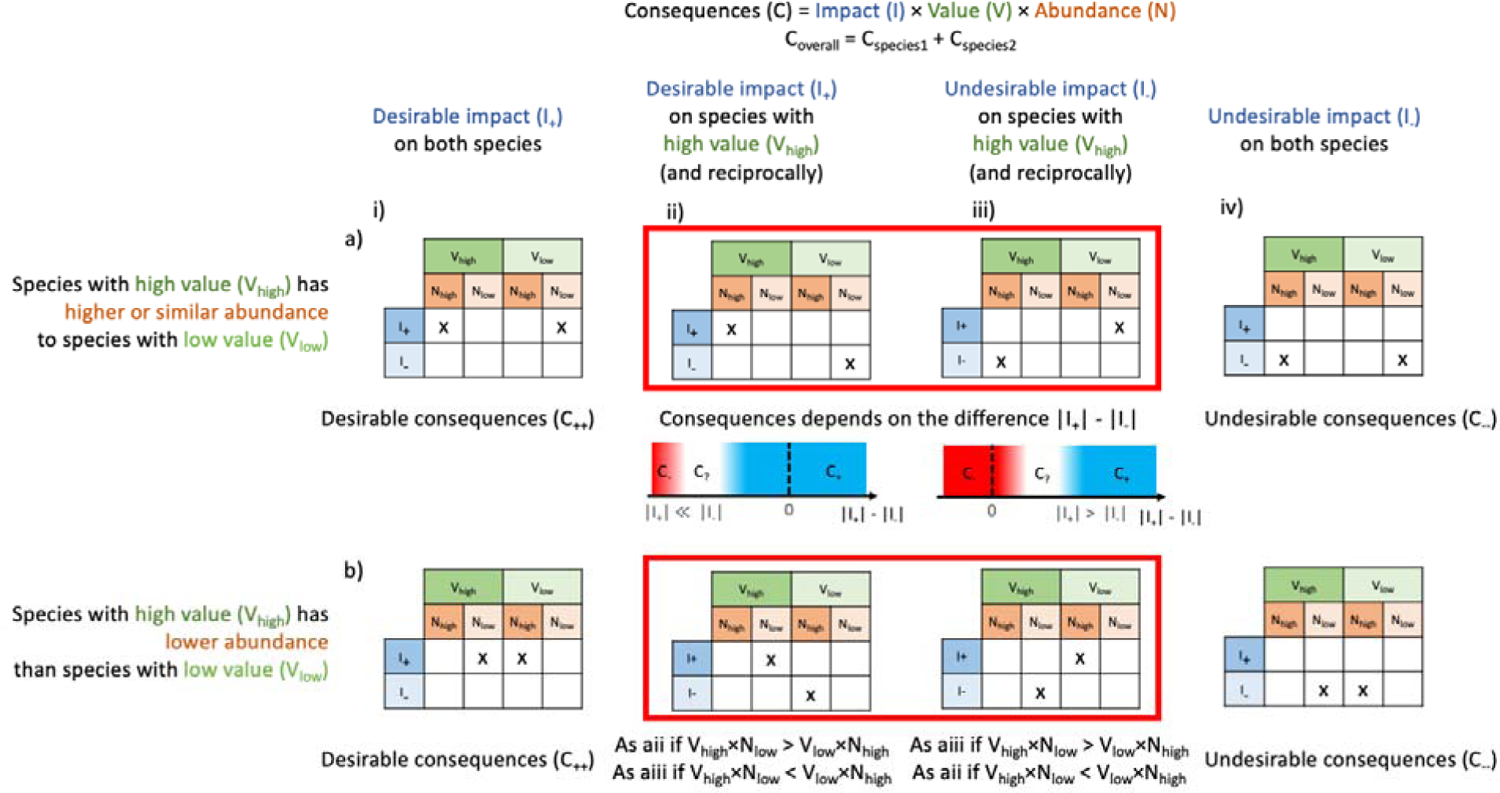
Applying the framework presented in Equation 1 to determine the likely consequence of a management action on a system with two species, highlighting possible moral dilemmas in red. In the case shown *a* is set to 1 for simplicity, but the two species have different inherent values V_high_ and V_low_ (i.e. how individuals are valued does not vary with abundance, but individuals of one species are valued more than the other species). The likely consequence changes with the relative abundance of the two species [top row (a) vs. bottom row (b)] and with whether the impact of the management intervention is positive (I_+_) or negative (I_-_) on the respective species [columns (i-iv)]. a) The species with high value has higher or similar abundance to the species with low value. If the impacts I_+_ and I_-_ have similar orders of magnitudes or |I_+_| > |I_-_|, aii generates positive consequences (C_+_) because V_high_ × N_high_ > V_low_ × N_low_. Similarly, if the impacts I_+_ and I_-_ have similar orders of magnitudes or |I_+_| < |I_-_|, aiii generates negative consequences (C-). If |I_+_| ≪ |I_-_| or |I_+_| > |I_-_| (for aii and aiii, respectively), the difference of impact can counter-balance V_high_ × N_high_ > V_low_ × N_low_, making desirable consequences undesirable and vice versa. However, the difference of magnitude between I_+_ and I_-_ at which this switch occurs is difficult to determine due to the different units of V, N, and I. This uncertainty corresponds to a moral dilemma due to a conflict between the desire to have a small positive impact for the species with the larger value and abundance, and the desire to avoid a very negative impact for the species with the lower value and abundance for aii. For aiii, the dilemma is due to a conflict between the desire to avoid a small negative impact for the species with the higher value and abundance, and the desire to have a very positive impact for the species with the lower value and abundance. b) The species with higher value V_high_ has the lower abundance N_low_. If impacts are different between the two species, the opposition between V and N will most likely generate moral dilemmas (C_?_). If V_high_ × N_low_ > V_low_ × N_high_, bii is equivalent to aii, and to aiii otherwise (and biii is equivalent to aiii, and to aii otherwise), but because value and abundance have different units, it is difficult to determine for which value and abundance V_high_ × N_low_ = V_low_ × N_high_. Therefore, an additional moral dilemma arises due to a conflict between the desire to avoid a negative impact for the larger population and the desire to avoid a negative impact for the species with the higher value.

**Table 2.** Set of questions to ask in order to evaluate Equation 1 and related concepts. The purpose is to guide users in exploring all the elements to consider when assessing the consequences of management actions rather than necessarily attempting a quantification of each. See Table 3 for factors to consider to answer these questions.

| Element of Equation 1 | Question | Mathematical formulation | Examples of interpretation |
| --- | --- | --- | --- |
| $V_s$ | What relative value do you place on individuals of different species? | What is the distribution of $V_s$? | - If a few species have a disproportionately high value compared to others, i.e. speciesism, the distribution of $V_s$ is highly skewed.<br>- If all species have a similar value, the distribution of $V_s$ is even. |
| $I_s$ | What measure(s) of impact do you consider? | What is the unit of $I_s$?<br>How to quantify $I_s$? | - If only individual survival matters, $I_s$ can be quantified as the probability of death, and assessed through surveys.<br>- If animal wellbeing matters, approaches based on physical aspect, stress, etc. can be used to quantify $I_s$. |
| $a$ | Do you value individuals or species? | What is the value of $a$? Is $a$ positive or negative? | - If individuals matter, $a > 0$, otherwise $a \leq 0$.<br>- If all individuals have the same value, $a = 1$. That means that common species will weigh more in the assessment of the conservation action. |
| | If you value species, should rare species have more values than common ones? | How negative is $a$? | - If all species must weigh the same, $a = 0$.<br>- If rare species should be given more importance, $a < 0$. |

If Equation 1 could be evaluated, for each measure of impact and each system of values, Equation 1 would produce relative rather than absolute values. The values of consequences *C* of a management action under different value systems and measure of impact cannot be directly compared with each other, because the unit and range of values of *C* can vary between value systems. Instead, Equation 1 can be used to rank a set of management actions for each value system or measure of impact based on their assessed consequences, to identify management actions representing consensus, compromises or conflicts amongst value systems.

Equation 1 is particularly useful to identify potential moral dilemmas, i.e. situations in which management options are conflicting under the same value system (Table 1). For example, if different types of impacts are considered simultaneously under a value system (e.g. economic vs cultural impacts, or lethal impacts vs. those causing suffering, see sections below), Equation 1 might rank management actions differently for these different impacts under the same system of moral values.

In some situations the implication of Equation 1 is clear. For example, if an impact is positive on a highly valued, highly abundant species, but slightly negative for a few individuals of another species that is not considered very important *(C = I_+_ x V_high_ x N_high_ + I_-_ x V_low_ x N_low_)*, the consequence will be positive (Figures 3aii, 4a). However, if the magnitude of the negative impact is much higher than that of the positive impact (|I_+_| ≪|I-|), the consequence can become negative. Similarly, if impact is negative for the species with the highest value and abundance, and positive for the other species *(C = I_+_ x V_high_ x N_high_ + I_-_ x V_low_ x N_low_)*, the situation is clear if positive and negative impacts have the same magnitude, but it will shift once the magnitude of the positive impact becomes higher than the magnitude of the negative impact (|I_+_| > |I-|; the difference of magnitude will likely be lower than in the first example, because of the differences in sign; Figures 3aiii, 4b). Since impact, value and abundance have different units, the thresholds at which these shifts occur are difficult to assess, and so the consequences can be highly debatable. This can create moral dilemmas, e.g. between the desire to have a small positive impact for a larger population with higher value and the desire to avoid a very negative impact for the species with the lower value and abundance (Figures 3aii, 4a); and between the desire to avoid a small negative impact for the larger population with the higher value and the desire to have a very positive impact for the species with the lower value and abundance (Figures 3aiii, 4b). Moral dilemmas will be even more likely to occur if the species with the higher value has the lower abundance *(C = I_+_ x V_high_ x N_high_ + I_-_ x V_low_ x N_high_* or *C = I_+_ x V_high_ x N_low_ + I_+_ x V_low_ x N_high_*; Figure 3bii,iii). If *V_high_ x N_low_ > V_low_ x N_high_*, the example depicted in Figure 3bii is equivalent to the example depicted in Figures 3aii, 4a described above, and Figure 3biii is equivalent to the example depicted in Figures 3aiii, 4b. If *V_high_ x N_low_ > V_low_ x N_high_*, the example depicted in Figure 3bii is equivalent to the example depicted in Figures 3aiii, 4b described above, and Figure 3biii is equivalent to the example depicted in Figures 3aii, 4a. As above, it is difficult to determine when the inequality will change direction because of the difference in the units of V and N. This reflects a moral dilemma due to a conflict between the desire to avoid a negative impact for the larger population and the desire to avoid a negative impact for the species with the higher value. In summary, uncertainty in the computation of Equation 1, and in particular the need to compare parameters with different units (i.e. impact, value, and abundance), can therefore be interpreted as a moral dilemma (Figures 3, 4).

In addition, some actions might not follow moral norms compared to others despite having more desirable consequences. For example, killing individuals may be considered less moral, but more efficient to preserve biodiversity or ecosystem services than using landscape management. Solving these moral dilemmas is complex, and beyond the scope of this publication, but approaches such as multi-criteria decision analyses (MCDA; Huang et al. 2011) may offer an avenue to do so (Goetghebeur and Wagner 2017).

Similarly, environmental conflicts will likely emerge when comparing the rankings generated by Equation 1 under different value systems considering different distributions of values, and different measures of impact. MCDA (Wittmer et al. 2006) and operational research (Kunsch et al. 2009), have also been proposed to resolve such conflicts. We nonetheless argue that, regardless of the capacity to resolve environmental conflicts (or moral dilemmas), being able to understand where these conflicts emerge from in Equation 1 can be beneficial for decision making.

In the following, we discuss the complexity of assessing the different variables and parameters of Equation 1 under different value systems using the set of primary questions defined above. By doing so, it becomes possible to identify ambiguity, difficulty of operationality, etc., to eventually move towards a good constructed attribute (although such a constructed attribute may not be reached in practice). We also discuss how, despite the difficulty to quantify the variables described above, this framework can be used as a heuristic (rather than operational) tool to capture the implications of considering different value systems for determining the appropriateness of a conservation action, and to better understand conservation disputes.

### NATURE DESPITE/FOR/AND PEOPLE

Over the past decade there has been some debate between proponents of traditional conservation, and those of new conservation (Table 1), as each group assumes different relationships between nature and people. Here, we show how the formal conceptualisation of Equation 1 could help clarifying the position of the new conservation approach in response to its criticisms (Kareiva 2014).

#### Nature despite people and traditional conservation

Traditional conservation is based on an ecocentric value system and seeks to maximize diversity of organisms, ecological complexity, and to enable evolution (Soulé 1985). For simplification, we will consider an extreme perspective of traditional conservation, championed by ‘fortress conservation’ (Siurua 2006, Büscher 2016), i.e. excluding humans from the moral community. To capture these aspects, consequences *C* in Equation 1 can be more specifically expressed as follows:

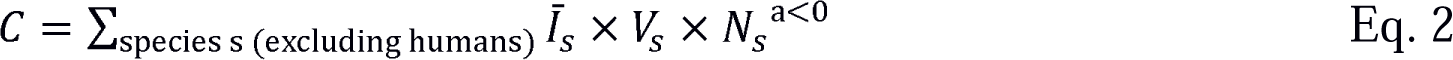

Assigning a stronger weight to rare species (*a* < 0) accounts for the fact that rare species are more likely to go extinct, decreasing the diversity of organisms. Evolution and ecological complexity are not explicitly accounted for in Equation 2. To do so, one may adapt Equation 2 and consider lineages or functional groups instead of species as the unit over which impacts are aggregated.

Because traditional conservation seeks to maximise diversity, *I_s_* can be defined as the probability of individuals dying. *I_s_* × *N_s_* will then be proportional to the extinction risk of a species (for an operational application, a proper model for extinction probability could be used in lieu of *I_s_* × *N*^a< 0^_s_). The *V_s_* distribution could be considered uniform over all species, in the absence of biases.

#### Nature for people and new conservation

New conservation considers that many stakeholders (“resource users”, Kareiva, 2014) tend to have an anthropocentric value system, and that conservation approaches that do not incorporate such a perspective will likely not succeed at maximizing diversity of organisms (Kareiva and Marvier 2012, Kareiva 2014). Under anthropocentrism, species are only conserved due to their utilitarian value, i.e. their effect on *I* for humans, rather than based on an inherent value *V*. Different groups of stakeholders are likely to be impacted differently (e.g. different monetary benefits / losses), and we propose the following extension of Equation 1 to account for this variability:

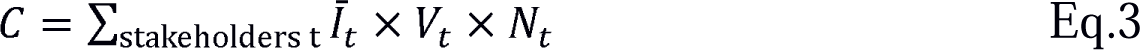

where I_t_ is the average impact of management on the group of stakeholders *t*, including indirect impacts through the effect of management of non-human species. \^ˉ^ can correspond to economic impacts, or encompass categorical measures of wellbeing (e.g. Bacher et al. 2018). *V_t_* is the value of the group of stakeholders *t*, and *N_t_* is its abundance (i.e. the number of people that compose it). Parameter *a* is set to 1, as this is considered to be an individual-based value system. Note that including inherent values *V_t_* in Equation 3 does not imply that we consider that different humans should be valued differently, but that is a view that some people have, and this needs to appear here to capture the full spectrum of perceived consequences of a management action.

New conservation holds an ambiguous perspective, stating that it should make “sure people benefit from conservation”, while at the same time does not “want to replace biological-diversity based conservation with a humanitarian movement” (Kareiva 2014). Using our framework, we interpret this to mean that one can design management actions that minimize consequences *C* under both Equations 2 and 3 (i.e. a mathematical expression of the convergence hypothesis; Norton 1986). Importantly, minimising Equation 3 is thereby a prerequisite for minimising impacts *I* and hence consequences *C* in Equation 2 (Figure 2). Under New conservation, Equation 2 can therefore be rewritten as follows:

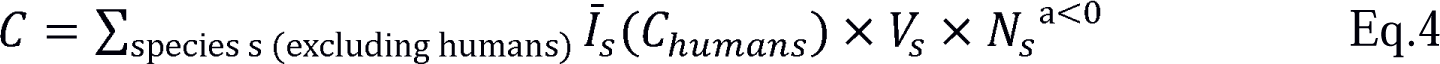

### Where *C_humans_* is computed using Equation 3

The link between biodiversity and ecosystem services is strongly supported, even if many unknowns remain (Chivian and Bernstein 2008, Cardinale et al. 2012), implying that high biodiversity can indeed support the provision of ecosystem services to humans. Such an approach will necessarily distinguish between “useful” species and others, and impacts will be perceived differently by different groups of stakeholders. Considering multiple types of impacts (economic benefits/losses, access to nature, health, etc.) while accounting for cultural differences, would increase the pool of useful species (comparing the resulting equation outputs using, for example, MCDA). The outcome of the two approaches would then potentially be more aligned with each other. This broad utilitarian perspective is captured in the most recent developments of new conservation approaches, which consider a wide range of nature contributions to people, rather than just ecosystem services (Díaz et al. 2018).

#### People and nature

People and nature views seek to simultaneously benefit human wellbeing and biodiversity (Figure 2). Under this perspective, Equations 2 and 3 should therefore be combined in a single approach, for example using MCDA (Huang et al. 2011; assuming these equations can indeed be operationally computed), to capture a more diverse set of value systems than Equations 2 and 3 alone, even if the two approaches generate divergent results.

We expressed traditional and new conservation with Equations 2, 3 and 4, which correspond to extreme interpretations of these two approaches (excluding humans or considering specific utilities of species). Doing so illustrates how our mathematical framework can capture in an explicit fashion the pitfalls of failing to explicitly define normative postulates for conservation approaches. As a result, Equations 2, 3 and 4 will likely generate conflicting results in the ranking of different management actions, especially if few types of impacts are considered. The debates over new conservation have taken place in a discursive fashion, which has not provided a clear answer to the values defended by this approach (Kareiva 2014, Soulé 2014, Doak et al. 2015). It has therefore been argued that the normative postulates of new conservation need to be more explicitly defined (Miller et al. 2011). Our framework could help doing so, by being more explicit about how new conservation would be defined relative to the traditional conservation and the people and nature perspective through the addition of specific terms to Equation 3 and a thorough comparison of the resulting equations. In particular, it would be interesting to explore, if inherent values are attributed to different species under a new conservation approach, how these values are determined compared to a traditional conservation approach (e.g. relational vs. intrinsic value; Chan et al. 2016; Table 1) and how their distributions differ.

### THE CASE OF ANIMAL WELFARE

The question of if and how animal welfare should be integrated into conservation practice is increasingly debated (Hampton and Hyndman 2018). Recently, conservation welfare (Table 1) has proposed to consider both the “fitness” (physical states) and “feelings” (mental experiences) of non-human individuals in conservation practice (Beausoleil et al. 2018). Based on virtue ethics rather than consequentialism, compassionate conservation (Table 1) also emphasises animal welfare and is based on the “growing recognition of the intrinsic value of conscious and sentient animals” (Wallach et al. 2018). It opposes the killing of sentient invasive alien species; the killing of sentient native predators threatening endangered species; or the killing of sentient individuals from a given population to fund broader conservation goals.

Despite the near-universal support of conservation practitioners and scientists for compassion towards wildlife and ensuring animal welfare (Russell et al. 2016, Hayward et al. 2019, Oommen et al. 2019), compassionate conservation has sparked vigorous responses (Hampton et al. 2018, Driscoll and Watson 2019, Hayward et al. 2019, Oommen et al. 2019, Griffin et al. 2020). Amongst the main criticisms of compassionate conservation is that the absence of action can result in (often well understood and predictable) detrimental effects and increased suffering for individuals of other or the same species (including humans), as a result of altered biotic interactions across multiple trophic levels, i.e. “not doing anything” is an active choice that has consequences (Table 3). However, since compassionate conservation is not based on consequentialism, it uses different criteria to assess the appropriateness of conservations actions (but see (Wallach et al. 2020) for responses to some criticisms). Our purpose here is not to discuss the relevance or irrelevance of virtue ethics for conservation (see (Griffin et al. 2020) for such criticism). Instead, we propose discussing animal welfare from the perspective of consequentialism (Hampton et al. 2018), i.e. more aligned with the approach of conservation welfare (Beausoleil et al. 2018), and to show how it may be aligned with or oppose the traditional and new conservation approaches.

**Table 3.** List of factors to consider regarding the effects of environmental management actions from an environmental ethics perspective.

| Factor | Influence on variables and outputs in Equations 1 to 5 |
| --- | --- |
| Biotic interactions | The impact or suffering of individuals from one species can be caused by individuals from another species, either through direct or indirect interactions. Management actions can therefore also have non-trivial indirect impacts on some species. |
| Capacity to provide ecosystem services | The presence of a specific species may increase the fitness/welfare of other species through the ecosystem services it provides. Since these effects can be difficult to quantify explicitly, the value of such species may be increased in Equations 1 to 4 to account for them. |
| Discounting rate | Rate at which impacts that occur in the future lose importance. |
| Impact quantification and commensurability | How the impacts of management actions are quantified is also dependent on value systems, as some impacts (such as death) may be considered incommensurable to others (such as suffering). |
| Responsibility from previous actions | Previous human actions on certain species, such as reintroduction of domesticated species or the introduction of alien species can change the perception of the public and therefore change the inherent value attributed to these species, or change the morality of an action, in addition to obviously having an impact on these species. |
| Spatial scale | The spatial scale will change the abundance $N$ and the number of species considered. As a result, a management action that is more beneficial than another at small scale may not be such at a larger scale, and reciprocally. Additionally, the spatial scale can change the inherent value of species, for example under nativism, or because of the range of cultures that are considered. |
| Temporal scale | The time frame over which the impact or the suffering of individuals is computed can change their values. Management actions may decrease welfare of individuals on the short term, but |
|  | be beneficial on the long term once the ecosystem has stabilised. Similarly, not culling some population may cause less suffering on the short term, but increase it in the future by disrupting ecosystem services, leading to population collapse due to lack of resources, etc. |
| Uncertainty of impact | The complexity of an ecological system can make the impact of management actions on different species difficult to assess precisely, therefore creating potential errors, especially in the presence of multiple biotic interactions. This may lead to an incorrect estimation of the consequences *C*. |
| Uncertainty of value expressions and preferences | Quantifying the value given by a person or a group of people to an individual is difficult, context-dependent, and highly subjective. Sensitivity analyses on the distribution of values can be used to account for such uncertainty. |

#### A mathematical conceptualisation of animal welfare

A consequentialist, sentientist perspective aims at maximizing happiness, or conversely minimising suffering, for all sentient beings, an approach also termed ‘utilitarianism’ (Singer 1980, Varner 2008). Suffering is therefore considered as a measure of impact (or, in mathematical terms, impact is a function of suffering, which can be expressed as *I*(*S_s_*) in Equation 1).

It has become widely accepted that animals experience emotions (de Waal 2011). Emotions have been shown to be linked to cognitive processes (Boissy and Lee 2014), which differ greatly among species (MacLean et al. 2012), and behavioural approaches have been used to evaluate and grade emotional responses (e.g. (Désiré et al. 2002); but see (Shriver 2006) and (Bermond et al. 2001) for different conclusions about the capacity of animals to experience suffering). We therefore postulate that the quantification of suffering is conceptually feasible in the context of the heuristic tool presented here. In a utilitarian approach, the inherent value of a species would therefore be a function of its capacity to experience emotions and suffering *E_s_*, which can be expressed as *V*(*E_s_*) instead of *V_s_* in Equation 1.

Under these considerations for defining impact and value of species, the consequences of a conservation action can be computed as a function of suffering of individuals from species *s S_s_*, their capacity to experience emotion and suffering *E_s_*, and the abundance of species *s*:

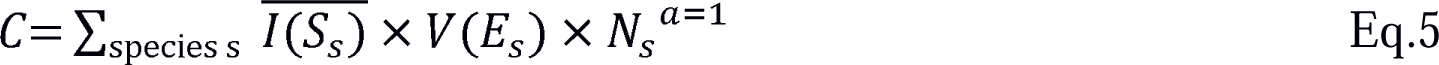

Although *V*(*E_s_*) should be measured in an objective fashion, many factors may influence the relationship between the inherent value and the emotional capacity of a species. For example, high empathy (Table 1) from the observer will tend to make the distribution uniform, whereas anthropomorphism and parochialism (Table 1) may lead to higher rating of the emotional capacities of species phylogenetically close to humans or with which humans are more often in contact, such as pets. Finally, we assumed that *a* = 1, to give equal importance to any individual regardless of the abundance of its species, as suffering and wellbeing are usually considered at the individual level (Beausoleil et al. 2018).

#### Assessing suffering in the presence and absence of conservation management actions

The short-term suffering resulting from pain and directly caused by lethal management actions, such as the use of poison to control invasive alien species (Twigg and Parker 2010) or the use of firearms and traps to cull native species threatening other native species (Proulx et al. 2016) or humans (Gibbs and Warren 2015), is the most straightforward type of suffering that can be assessed, and is usually sought to be minimised in all conservation approaches. Suffering can have many other causes, and suffering of an individual may be assessed through a wide variety of proxies, including access to food and water, death, number of dead kin for social animals, physiological measurements of stress hormones, etc. Suffering can take various forms, and commensurability can be an issue (Table 3), making the distinction between the morality of lethal actions and non-lethal suffering complex. Non-lethal suffering can result from unfavourable environmental conditions (e.g. leading to food deprivation) and occur over long periods, while lethal actions could be carried out in a quick, non-painful fashion (Shao et al. 2018), and even lead to improved animal welfare (Wilson and Edwards 2019), but may be deemed immoral.

The concept of animal welfare and how to measure it is extremely complex (Beausoleil et al. 2018), and defining it precisely is beyond the scope of this study. We nonetheless advocate a conceptual approach that takes into account indirect consequences of management actions within a certain timeframe and consider uncertainty (Table 3). Direct and indirect biotic interactions may be explicitly modelled to quantify the impact on animals and their suffering. Simulation models can also make projections on how populations may change in time, accounting for future suffering.

#### Are traditional conservation and animal welfare compatible?

It has been argued that sentientism and ecocentrism are not fully incompatible (Varner 2011). The relationship between biodiversity and animal suffering can be formalised more clearly using the traditional conservation and the sentientist Equations 2 and 4, to explore if the same management action can minimize the consequences evaluated using the two equations (see also Appendix S2 for the application of the framework to theoretical cases). The main difference with the traditional vs new conservation debate here is that Equations 2 and 4 share a number of species, whereas the new conservation Equation 3 only contains humans, which are excluded from Equation 2. Even though the variables of Equation 4 differ from those of Equation 2 (*V* and *I* are computed differently, and the value of *a* is different), it is possible that these equations will vary in similar way for different management actions due to their similar structure, although this would depend on the variety of impacts on humans that are considered in Equation 3. Finally, as for the people and nature approach, the consequences of sentientist and ecocentric approaches can be evaluated in combination, as suggested by conservation welfare (Beausoleil et al. 2018), using tools such as MCDA (Wittmer et al. 2006, Huang et al. 2011).

One issue that may be irreconcilable between ecocentric approaches such as traditional conservation and approaches based on sentientism is the fate of rare and endangered species with limited or no sentience. Under utilitarian sentientism, the conservation of non-sentient species ranks lower than the conservation of sentient species, and consequently they are not included in Equation 4. For example, endangered plant species that are not a resource for the maintenance of sentient populations would receive no attention, as there would be few arguments for their conservation. Traditional conservation would focus on their conservation, as they would have a disproportionate impact in Equation 2, due to low abundance leading to a high value for *N ^a<0^*.

Finally, it is important to note that the current body of knowledge shows that the link between biodiversity and animal welfare mentioned above especially applies to the increase of native biodiversity. Local increase of biodiversity due to the introduction of alien species may only be temporary due to extinction debt (Kuussaari et al. 2009) and often results in a reduction of ecosystem functioning (Cardinale et al. 2012). Therefore, it is important to distinguish between nativism (Table 1) and the detrimental effects of *invasive* alien species on biodiversity and ecosystem functioning and services (Bellard et al. 2016). Nativism would result in increasing the inherent value *V_s_* of native species (Figure 1), whereas in the second case, insights from science on the impact of invasive alien species would modify the distribution *I*(*S*) rather than the distribution *V_s_*. This can also apply to native species whose impacts on other species, such as predation, are increased through environmental changes (Carey et al. 2012).

### UNRESOLVED QUESTIONS AND LIMITATIONS

From an operational perspective, this framework shares similarities with mathematical approaches used in conservation triage (Bottrill et al. 2008), but has two crucial differences. First, conservation triage is an ecocentric perspective with variables that are comparatively easy to quantify. Bottrill et al. (2008) provided an example using phylogenetic diversity as a measure of value *V*, and a binomial value *b* to quantify biodiversity benefit that can be interpreted as the presence or absence of a species (i.e. *I* = 1 / b). Because it is ecocentric, local species abundance is not considered, which corresponds to setting *a* = 0. In this example, consequences (*C*) in the general Equation 1 are therefore defined simply by *V* / *b*.

In contrast, our framework allows more flexibility to encompass a range of value systems, as shown above. However, given that the data needed for quantifying parameters of Equations 1 to 4 related to value, impact, emotional capacity and suffering are scarce and often very difficult to measure, this framework in its current form would be difficult to use as a quantitative decision tool to evaluate alternative management actions, contrary to triage equations. Rather, our equations decompose the question underlying many controversies around management decisions in conservation: what or who is valued, how, and to what extent?

Despite the difficulty to apply the framework, it can guide the search for approaches that may be used to develop quantification schemes for the different parameters of the framework and therefore obtain a better appreciation of the different facets of the valuation of nature. For example, grading systems may be developed to assess impact and suffering based on various indicators, including appearance, physiology, and behaviour (Broom 1988, Beausoleil et al. 2018). For assessing the value of different species, questionnaires may be used to assess how different species are valued by people, and influenced by their social and cultural background, similar to what has been done to assess species charisma (Colléony et al. 2017, Albert et al. 2018). It will nonetheless be important to acknowledge the corresponding uncertainties in the assessment of impact and value, differences in perception among societal groups for different taxa and potential shifts in perception over time (Table 3).

The second difference from conservation triage is that the latter considers additional criteria that were not addressed here, including feasibility, cost, and efficiency (including related uncertainties). The combination of these different perspectives calls for appropriate methods to include them all in decision making, which can be done using MCDA (Huang et al. 2011). Here, good communication and transparency of the decision process is key to achieve the highest possible acceptance across stakeholders, and to avoid biases in public perception (see case studies below for examples).

The issue of spatial and temporal scale also warrants consideration (Table 3). In the case of a species that may be detrimental to others in a given location but in decline globally, the spatial scale and the population considered for evaluating the terms of Equations 1 to 4 is crucial to determine appropriate management actions. Similarly, management actions may also result in a temporary decrease in welfare conditions for animals, which may increase later on (Ohl and Van der Staay 2012), or the impacts may be manifested with a temporal lag. In that case, determining the appropriate time period over which to evaluate the terms of Equations 1 to 4 will not be straightforward. Impacts will be of different importance depending on whether they occur in the short- or long-term, especially since long-term impacts are harder to predict and involve higher uncertainty. Discount rates (Table 3) may therefore be applied, in a similar way they are applied to the future effects of climate change and carbon emissions (Essl et al. 2018), or to assess the impact of alien species (Essl et al. 2017).

Equations 1 to 5 assume that all individuals from a given species have the same value or emotional capacities (or use the average of the value across individuals). However, intraspecific differences in value may be important for conservation. For example, reproductively active individuals contributing to population growth/recovery may be given a higher value in an ecocentric perspective. Trophy hunters might prefer to hunt adult male deer with large antlers. Intraspecific value may also vary spatially, for example between individuals in nature reserves or in highly disturbed ecosystems. Equation 1 may therefore theoretically be adapted to use custom groups of individuals with specific values within species, similar to Equation 3.

Finally, it is crucial to account for biotic interactions in our framework to comprehensively assess the indirect impacts of management actions on different species (Table 3). Some species with low values *V_s_* in a certain value system may be crucial for assessing the impact *I_s_* on other species. These biotic interactions will therefore determine the time frame over which the framework should be applied, as impacts on one species at a given time may have important repercussions in the future. These biotic interactions can be complex, and several tools, such as simulation models and ecological network analyses can be used to capture them. Concepts such as keystone species (Mills et al. 1993) can also offer a convenient way to overcome such complexity by modifying *V_s_* rather than *I^-^_s_*. Let us assume that a management action will have a direct impact on a keystone species, which will result in indirect impacts on multiple other species with inherent values. Increasing the value of the keystone species can result in the same assessment of *C* as to explicitly model the biotic interactions and compute the resulting indirect impacts *I^-^_s_*.

### CASE STUDIES ILLUSTRATING ETHICAL CONFLICTS IN CONSERVATION DECISIONS

In the following, we present three case studies where conservation actions have either failed, had adverse effects, or were controversial, and we explore how our framework can help to identify normative postulates underlying these situations. Although these case studies have been discussed at length in the articles and reports we cite, we argue that our framework helps capture the different components of the controversies in a more straightforward and objective fashion than using a discursive approach that might require either emotionally loaded language or more neutral but less understood neologisms.

#### Invasive alien species management: the case of the alien grey squirrel in Italy

The grey squirrel (*Sciurus carolinensis*) is native to North America and was introduced in various locations in Europe during the late nineteenth and the twentieth century (Bertolino 2008). It threatens native European red squirrel (*Sciurus vulgaris*) populations through competitive exclusion and as a vector of transmission of squirrel poxvirus in Great Britain (Schuchert et al. 2014). Furthermore, it has wider impacts on woodlands and plantations, reducing value of tree crops, and potentially affects bird populations through nest predation (Bertolino 2008).

Based on the impacts of the grey squirrel, an eradication campaign was implemented in 1997 in Italy, with encouraging preliminary results (Genovesi and Bertolino 2001). However, this eradication campaign was halted by public pressure from animal rights movements. The strategy of the animal rights activists consisted in (i) humanising the grey squirrel and using emotive messages (referring to grey squirrels as “Cip and Ciop”, the Italian names of the Walt Disney “Chip and Dale” characters) and (ii) minimising or denying the effect of grey squirrel on native taxa, especially the red squirrel (Genovesi and Bertolino 2001). In addition, the activists did not mention (iii) the difference in abundance between a small founding population of grey squirrels that could be eradicated by managers, and a large population of native red squirrels that would be extirpated or severely impacted by grey squirrels if control was not implemented.

Genovesi & Bertolino (2001) explain that the main reason for the failure of the species management was a different perspective on primary values. The conservation managers, favouring eradication, based their decision on species valuation, following traditional conservation. The animal rights activists, opposed to control, focussed on animal welfare. Applying the framework, and assuming an individual-based value system (*a* = 1 in Equation 1), three questions are apparent (Table 2):

i. Are the values of red and grey squirrels different?
ii. What types of impact are we considering?
iii. Is the population of red squirrels impacted by grey squirrels larger than the population of grey squirrels to be controlled?

The arguments of animal rights activists led to the following answers to these three questions. (i) The humanisation of the grey squirrel consists of increasing the perception of its emotional capacity *E_gs_* > *E_rs_* (and therefore *V*(*E_gs_*) > *V*(*E_rs_*)). (ii) Minimising the impact of the grey squirrel is equal to restricting the time scale to a short one and to likely minimising the amount of suffering *S* caused by grey squirrels on other species (under a sentientist perspective), or the number of red squirrels that will die because of grey squirrels (under a biocentric perspective). In other words, *S_gs_* = *S_rs_* (and therefore *I*(*S_gs_*) = *I*(*S_rs_*)) or *I_gs_* = *I_rs_* without management and *S_gs_* > *S_rs_* (and therefore *I*(*S_gs_*) > *I*(*S_rs_*)) or *I_gs_* > *I_rs_* under management. (iii) Not mentioning differences in species abundance implies that the impacted populations of red and grey squirrels would have the same size under any management. Following these three points, the consequences under management *C_m_ = I(S_gs_) × V(E_gs_) + I(S_rs_) × V(E_rs_)* are higher than without management, due to the increase in *V*(*E_gs_*) and *I*(*S_gs_*). The application of our framework therefore allows to clarify a discourse whose perception could otherwise be altered because of techniques such as appeal to emotion.

The framework can thus be used to provide recommendations for what the advocates for the eradication campaign would have needed to have done: i) increase the value *E_rs_* of red squirrels in a similar way as what was done for grey squirrels, so that their relative values compared to grey squirrels would remain the same as before the communication campaign by the animal right activists; ii) better explain the differences in animal death and suffering caused by the long-term presence of the grey squirrel compared to the short-term, carefully designed euthanasia protocol, which would avoid a subjective perception of the distribution of *S*; and iii) highlight the differences in the number of individuals affected. The consequences would then be computed as *C = V(E_gs_) x I(S_gs_) x N_gs._+ V(E_rs_) x I(S_rs_) x N_rs_*. In that case, assuming for simplification the same suffering through euthanasia for grey squirrels as red squirrels suffer from the grey squirrels, and the same value to individuals of each species (i.e. avoiding nativism), the mere differences N_rs_ > N_gs_ in abundance would lead to a higher value of *C* without management. This would further increase by extending the impacts of grey squirrels to other species, as mentioned above.

A more fundamental issue, however, is that in some value systems it would not be acceptable to actively kill individuals, even if that meant letting grey squirrels eliminate red squirrels over long periods of time (Wallach et al. 2018). The reluctance to support indirectly positive conservation programs is a common issue (Courchamp et al. 2017). Whether an acceptable threshold on consequences over which killing individuals could be determined through discussion would depend, in part, on the willingness of the affected parties to compromise.

#### De-domestication: the case of Oostvaardersplassen nature reserve

De-domestication, the intentional reintroduction of domesticated species to the wild, is a recent practice in conservation that raises new ethical questions related to the unique status of these species (Gamborg et al. 2010). Oostvaardersplassen is a Dutch nature reserve. Reserve managers, recognising that grazing by large herbivore was a key natural ecosystem process that had been lost, decided between 1983 and 1992 to reintroduce red deer (*Cervus elaphus*), and two domesticated species (Heck cattle, *Bos primigenius*, and konik horses, *Equus ferus caballus*) (ICMO2 2010). The populations of these three species increased rapidly, as natural predators were missing and, as a result of a ‘non-intervention-strategy’, no active population control measures were implemented. The project was widely criticized when a considerable number of individuals died from starvation during a harsh winter, resulting in the subsequent introduction of culls.

From a traditional conservation perspective, disregarding animal welfare and focusing on species diversity and ecological restoration, the project was a success. The introduction of the three herbivore species led to sustainable populations (despite high winter mortality events), and ensured stability of bird populations without the need for further interventions (ICMO2 2010), i.e. the conditions of many species were improved (the impact was lowered), leading to improved consequences *C* for biodiversity overall (Equation 2). In other words, since more individuals from all species survived (*I* increased in Equation 2), *C* improved overall, regardless of differences in value or abundance between species (a multi-species generalisation of the Figure 2i).

However, the welfare of individuals from the three charismatic large herbivorous species became a point of conflict. In terms of the framework, it appears that the conflict was driven by considering the outcome of Equation 5 in addition to that of Equation 2 to estimate the overall evaluation of the management approach, i.e. a change from only considering impacts on individual survival to also considering impacts based on suffering, with the acknowledgement that *E_s_* should be considered (Ohl and Van der Staay 2012). Not considering Equation 5 would mean that *C* = 0 under sentientism, but acknowledging the existence of *E_s_* implies that *C = V(E_s_) x I(S_s_) x N_s_^1^* VI becomes non-null. Changes in perspective over time should therefore be taken into account when implementing conservation management actions, and adaptive management approaches should be considered. A possible explanation for this shift in attitude is the notion of responsibility (Table 3). Culling animals might be acceptable in some cases, but might not be if these individuals were purposefully introduced, which may lead to considering a sentientist perspective.

The reserve managers have examined a number of sustainable measures to improve the welfare of individuals from the three species (therefore decreasing *S_s_* to compensate the increase in *V_s_*). Among those were recommendations to increase access to natural shelter in neighbouring areas of woodland or forestry, to create shelter ridges to increase survival in winter as an ethical and sustainable solution, and to use early culling to regulate populations and avoid suffering from starvation in winter (ICMO2 2010). This example shows how a combination of two complementary management actions (the rewilding of the OVP and the provision of shelter) led to minimised consequences under both the traditional conservation and the sentientist Equations 2 and 5, whereas only rewilding would increase consequences under Equation 5. Culling may still face opposition based on moral arguments though. Interestingly other approaches, such as the reintroduction of large predators, were also considered but discarded due to a lack of experience and too many uncertainties in efficiency (ICMO2 2010). Our suggested framework could be adapted to explore the consequences of culling vs. increased mortality through the reintroduction of large predators, noting again that some stakeholders may make moral distinctions between natural mortality and human-induced mortality.

#### Trophy hunting

Trophy hunting, the use of charismatic species for hunting activities, has been argued to be good for conservation when revenues are reinvested properly into nature protection and redistributed across local communities, but faces criticisms for moral reasons (Lindsey et al. 2007b, Di Minin et al. 2016). The action of killing some individuals to save others might be incompatible with a deontological perspective, but, assuming a consequentialist perspective, the framework can be applied to formalise the assessment of different management options. Note that here, we are not considering the ethics of how the hunt itself is carried our (e.g. canned hunting vs. a “fair chase”) nor how animals are reared (i.e. whether they can express their natural behaviours), recognising that both these factors would need to be considered when making a decision.

In traditional conservation, trophy hunting is desirable if it directly contributes to the maintenance of species diversity. That is, it should decrease impacts *I* evaluated as individual survival over all or the majority of species with high inherent value, leading to improved consequences for biodiversity *C* in Equation 2 (a multi-species generalisation of Figures 2i and 2ii). The potential of trophy hunting to contribute to the maintenance of biodiversity is via creating economic revenues, i.e. an anthropocentric perspective, and it therefore falls under the umbrella of new conservation (Figure 2; Equation 4). In theory, trophy hunting should lead to lower consequences than doing nothing for both the traditional and new conservation (Equations 2, 3 and 4), and therefore for the ‘people and nature’ approach, as they are in this case not independent from each other (Lindsey et al. 2007a). Many social and biological factors currently affect the efficacy of trophy hunting as a conservation tool. Corruption and privatisation of the benefits have sometimes prevented the revenues to be reinvested into conservation, but also to be redistributed across local communities, whereas doing so has been shown to increase their participation in conservation actions with proven benefits for local biodiversity (Di Minin et al. 2016). In other words, a decrease in the anthropocentric Equation 2 leads to a decrease in the ecocentric Equation 3, but the causal link (Equation 4) is still supposed to be valid. In addition, trophy hunting can lead to unexpected evolutionary consequences (Coltman et al. 2003), overharvesting of young males (Lindsey et al. 2007b), and disproportionate pressure on threatened species (Palazy et al. 2011, 2012, 2013) and therefore to population declines and potential detrimental effects on biodiversity. That means that I(C_humans_) in Equation 4 should be carefully examined. Despite these issues, it has been argued that banning trophy hunting may create replacement activities that would be more detrimental to biodiversity (Di Minin et al. 2016).

From an animal welfare perspective, trophy hunting appears to be in direct contradiction with a decrease in animal suffering, and has been criticised by proponents of compassionate conservation (Wallach et al. 2018). However, as for the culling of invasive alien species, we suspect the story is more complex. First, there may be direct benefits for animal welfare, if money from trophy hunting is reinvested in protection measures against poaching (if such poaching causes, on balance, more suffering). Second, to our knowledge, only few studies have compared the welfare of individual animals to quantify the elements of the sentientist Equation 5 (for example assessed through access to resources) in areas where trophy hunting is practiced and where it is not. Given the links between biodiversity and animal welfare described above, it seems plausible that good practice in trophy hunting may benefit the welfare of individuals from other and from the same species.

## CONCLUSIONS

A variety of value systems exist in conservation, which are based on different underlying normative postulates and can differ between stakeholders, resulting in differing preferences for conservation practices among people. Here, we have proposed a framework with a formal set of equations to conceptualize and decompose these different perspectives from a consequentialist point of view. In this framework, the different value systems supported by different conservation approaches follow the same structure, but can differ in the variables that are used, and in the values they are taking. Such a formalisation by necessity does not capture the full range of complex and nuanced real-world situations in environmental decision-making, and the elements of the equations can be difficult to estimate. However, this framework is not intended to be an operational approach readily applicable across all value systems. Rather, the mathematical structure and the systematic examination of the elements of the framework provides a method to make their underlying value systems and the resulting conflicts explicit and transparent, which is essential for the planning and implementation of pro-active management. The search for consensus in conservation can be counter-productive and favour status-quo or ‘do nothing’ against pro-active management (Peterson et al. 2005), however our framework may help identify hidden commonalities between seemingly antagonistic stances. We hope that this framework can foster fruitful debates and thus facilitate the resolution of contested conservation issues, and will ultimately contribute to a broader appreciation of different viewpoints. In an increasingly complex world shaped by human activities, this is becoming ever more important.

## Acknowledgements

We thank Franck Courchamp, Vincent Devictor, Jordan Hampton, Jonathan Jeschke, and Thomas Potthast for extremely useful comments on previous versions of this manuscript. This research was funded through the 2017-2018 Belmont Forum and BiodivERsA joint call for research proposals, under the BiodivScen ERA-Net COFUND programme, and with the funding organisations Austrian Science Foundation FWF for GL, BL, AS, FE, and SD (BiodivERsA-Belmont Forum Project ‘Alien Scenarios’, grant no. I 4011-B32; grant no. I3757-B29). AP was funded by Conicyt PIA CCTE AFB170008 and ANID PIA FB210006. IJ acknowledges support by the J. E. Purkyně Fellowship of the Czech Academy of Sciences. JRUW thanks the South African Department of Forestry, Fisheries, and the Environment (DFFE) for funding noting that this publication does not necessarily represent the views or opinions of DFFE or its employees. We appreciate the helpful comments of Tina Heger and one anonymous reviewer.

